# Detecting DNA Methylation using the Oxford Nanopore Technologies MinION sequencer

**DOI:** 10.1101/047142

**Authors:** Jared T Simpson, Rachael Workman, P. C. Zuzarte, Matei David, L. J. Dursi, Winston Timp

## Abstract

Nanopore sequencing instruments measure the change in electric current caused by DNA transiting through the pore. In experimental and prototype nanopore sequencing devices it has been shown that the electrolytic current signals are sensitive to base modifications, such as 5-methylcytosine. Here we quantify the strength of this effect for the Oxford Nanopore Technologies MinION sequencer. Using synthetically methylated DNA we are able to train a hidden Markov model to distinguish 5-methylcytosine from unmethylated cytosine in DNA. We demonstrate by sequencing natural human DNA, without any special library preparation, that global patterns of methylation can be detected from low-coverage sequencing and that the methylation status of CpG islands can be reliably predicted from single MinION reads. Our trained model and prediction software is open source and freely available to the community under the MIT license.

## Main Text

Epigenetics is the study of heritable changes in phenotype through a mechanism other than genetic sequence, e.g., DNA cytosine methylation, histone post-translational modifications, chromatin compaction and nuclear organization. The epigenome acts as a cellular memory, retaining information after cell division. The epigenetic state of the cell is critical for regulating gene expression and cellular response to stimuli; it provides a mechanism for the cell to retain information even through multiple generations. Recent projects including the Encyclopedia of DNA Elements^1^ and the Epigenetic Roadmap^2^ sought to characterize the epigenome. Epigenetic signatures differ between somatic cells and stem cells^3^, between different tissue types^4^, and between normal and cancer samples^4^. Many, if not all, human diseases are a result of erroneous gene regulation, through mechanisms of genetic mutation, altered signal transduction, or epigenetic alteration.

The epigenetic state is highly mutable, changing more easily and rapidly than genetic mutation or alteration. This results in epigenetic variation among genetically homogeneous populations, a variation that can grant selective advantage to cells or organisms. This is especially apparent when comparing cancer and normal tissue – though normal tissue has clearly defined epigenetic signatures, cancer epigenetic signatures vary from sample to sample^4^, and even from cell to cell^5^. With the advent of high throughput DNA-sequencing technologies, methods have been developed for examining nuclear organization^6^, chromatin state/histone post translational modifications^7^, chromatin accessibility^8^ and methylation state^9^. Currently, 5-methylcytosine is typically profiled by treating DNA with sodium bisulfite to convert unmethylated cytosine to uracil which is subsequently read as thymine when short read sequenced. This widespread technique is limited in that it requires a separate assay, results in extensive DNA fragmentation^10^ resulting in material loss and does not reveal long-range single-read patterns of methylation due to the short read length. The Pacific Biosciences RSII instrument is able to directly detect methylation, without chemical treatment of the DNA prior to sequencing, at both known and novel motifs with long reads via polymerase kinetics^11^.

In previous work it has been shown that 5-methylcytosine can be distinguished from cytosine by careful analysis of the electric current signals that are measured by nanopore-based sequencing devices^12,13^. A nanopore based DNA sequencer, the Oxford Nanopore Technologies MinION, is now commercially available. In this paper we develop and demonstrate a method to directly detect DNA modifications, in this case 5-methylcytosine, based only on the electrical readout of the MinION without any chemical treatment. We characterize the error rate of our approach by sequencing negative and positive control samples and assess the biological context of the methylation patterns we detect using low-coverage human genome sequencing data.

## Results

Our results have two main sections. First, we describe the development and training of a probabilistic model of the electric signals measured by the nanopore for DNA containing 5-methylcytosine in a CpG context using methylated *E. coli* DNA. Second, we apply this trained model to detect 5-mC using MinION reads of genomic DNA from a human cell line. We compare the detected methylation patterns to sequenced negative and positive controls consisting of PCR-treated (to remove methylation) and enzymatically methylated (to methylate all CpGs) DNA for the same sample.

Throughout the results we will describe the data sets with a structured name containing the method used to prepare the DNA, the lab that sequenced the sample and the date of sequencing. For example data set PCR-timp-021216 refers to PCR-amplified DNA sequenced by the Timp lab on February 12th, 2016. For a complete description of the data sets we generated for the study see **Supplemental Table 5**.

## Model Development and Training

In previous work we and others have demonstrated how hidden Markov models can be used to analyze nanopore sequencing data ^14–17^. In our HMM we compute the probability of observing a sequence of signal-level *events* measured by the MinION given a nucleotide sequence *S* (see the **Supplementary Note**). In this work we extend our hidden Markov model used for de novo assembly and SNP calling ^14,18^ to detect methylation.

A hidden Markov model contains a set of emission distribution functions that model the probability of some observed data for each state of the HMM. For sequencing MinION data the observations are segmented electric current samples which are termed *events*. Each event represents a current level measured in picoamps over some interval of time. The emission distribution functions of our HMM (and others’) model the event observations using DNA *k*-mers (in this work *k*=6). The emission functions are Gaussian distributions with mean and standard deviation that depends on the sequence of the particular *k*-mer. For example the probability of observing event *e*_*i*_ given that it came from *k*-mer *k* is:

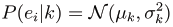

**Figure 1a** shows an example of events, and the corresponding Gaussian distribution, for the 6-mer AGGTAG.

**Figure 1:**
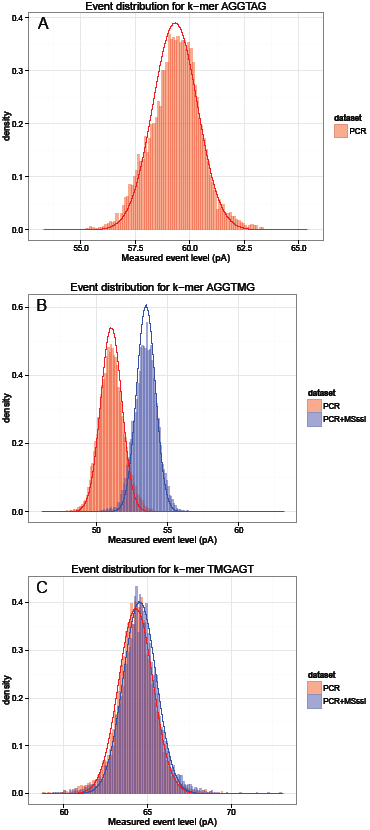
Differences between methylated and unmethylated 6-mers. Panel A is a histogram of the measured event current levels aligned to 6-mer **`AGGTAG`** for E. coli PCR sample (timp-021216). The solid red line is the Gaussian distribution fit to these observations. In panel B the measured events and Gaussian fits are shown for 6-mer **`AGGTMG`** for a methylated (blue) and unmethy-lated (red) data set. In panel C are the events and Gaussian fit for 6-mer **`TMGAGT`** for the same two data sets. In all panels the events shown are from the template strand.

Oxford Nanopore provides a reference set of parameters for all 4,096 6-mers over the standard nucleotide alphabet (A,C,G,T) for each of the three “strand models”. The first strand model is called the “template” model and is used for the first strand of DNA that passes through the pore. The other two models are for the “complement” strand that is sequenced after the template strand when a hairpin adaptor is used to make a “two-direction” (2D) read. The two complement models are called “complement.pop1” and “complement.pop2”. The Metrichor basecaller selects one of the two complement models depending on whether it inferred that double-stranded DNA reformed as the complement strand passed through the pore.

We developed a procedure to learn new parameters 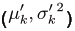 for each *k*c-mer for each of the three strand models by aligning nanopore reads to a known reference genome. This training method is described in **Supplementary Note - Training Emission Distributions**.

To test and validate our training procedure we used nanopore reads from PCR-amplified DNA from two *Escherichia coli* strains, one sequenced sample of K12 MG1655 and two sequenced samples of K12 ER2925. The PCR amplicons are generated without defect or methylation marks, so we expect that events measured by the MinION, and hence our trained parameters, will closely match the ONT reference parameters. To assess this we trained new parameters for the three PCR data sets independently and counted the number of *k*c-mers whose trained means differed from the ONT reference value by more than 0.1 pA, 0.5pA, 1.0pA and 2.0pA for each of the three strand models. These results are summarized in **Supplementary Table 1**. For all three PCR samples the trained template and complement.pop1 parameters closely match the values expected from the ONT reference models? very few *k*-mers have a trained mean that is more than 1pA different than the ONT mean. However, for one of the samples (PCR-timp-113015) the complement.pop2 model has many *k*-mers with a trained mean more than 1pA different than the ONT value. This result was explained by a low flowcell temperature of 31-32C, relative to the expected 37C, which Oxford Nanopore Technologies indicated might affect the signal levels. We resequenced this sample after eliminating the temperature variation by controlling room ventilation more precisely to obtain the dataset PCR-timp-021216. The trained parameters for this run are consistent with the ONT reference models for all three strands.

We then enzymatically methylated the same two PCR-amplified ER2925 DNA samples from above with M.SssI methyltransferase (Zymo), which converts cytosine in a CpG context to 5-mC. We validated the methylation state of these samples using whole genome bisulfite sequencing on an Illumina MiSeq, resulting in 89% and 97% methylation for PCR+M.SssI-timp-113015 and PCR+M.SssI-timp-021216. Each sample was run on its own flowcell and we ran our training procedure for these runs as before. In this case for both runs and all three strand models we observe many *k*-mers with trained means that are more than 1pA different when compared to the ONT reference models (**Supplementary Table 2**). As these differences are consistent across the two runs and the three strand models, and far exceed the difference we observed for the anomalous sample (PCR-timp-113015 complement.pop2), we conclude that the differences are caused by 5-mC.

Next, we sought to learn a new set of strand models for a *k*-mer collection that includes methylated cytosine. We refer to this alphabet as the CpG alphabet which contains the symbols (A,C,G,T,M) where M stands for 5-mC and can only appear in a CpG context. We trained new parameters over this expanded alphabet by converting all CG dinucleotides in the *E. coli* reference genome to MG. The results of training using the PCR + M.SssI data set (timp-021216) over this expanded alphabet are presented in (**Supplementary Table 4)**. Again, we observed many *k*-mers whose trained means are more than 1pA different than the ONT reference parameters, for all three strand models (in this case we compared the mean for methylated *k*-mers to the unmethylated version in the ONT reference model).

To confirm that this effect was not due to simply increasing the alphabet size of the model, which implicitly trains 7-mers rather than 6-mers for a subset of *k*-mers, we ran the same procedure on the PCR (unmethylated) datasets in parallel. Here we do not observe the same magnitude of difference for the trained means (**Supplementary Table 3**). **Figure 1b** is an example of a strong difference in signal for 6-mer AGGTMG, between the unmethylated data set PCR-timp-021216 and the methylated data set PCR+M.SssI-timp-021216. We note however that not all 6-mers demonstrate a difference in signal (**Figure 1c**).

We determined that the difference in trained mean when compared to the ONT reference model is dependent on the position of the methylated base in the 6-mer (**Figure 2**). When the methylated base is in the 5th and 6th position of the *k*-mer (*k*-mers with pattern abcd**M**f and abcde**M**) we see a consistent shift towards higher current observations. However, when the methylated base is in the first position of the *k*-mer (pattern **M**bcdef) very little difference with respect to the ONT reference model is observed.

**Figure 2:**
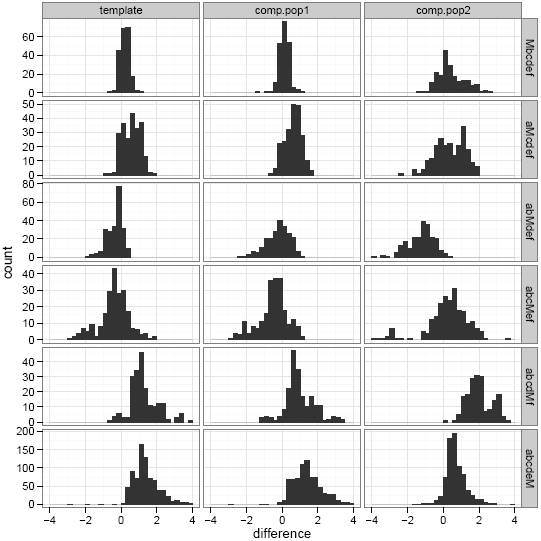
Position dependence of methylation parameter shift. A histogram of the difference between the trained mean for the methylated (PCR+M.SssI-timp-021216) dataset and the ONT reference model. Each pane is a set of *k*-mers from one of the three strand models (columns). Each row is the subset of *k*-mers with the methylated base in the first position (Mbcdef), the second position (aMcdef), and so on. We only included *k*-mers that contained a single methylation position. We restrict the plotting range to differences in the range −4 to 4 so some outliers are not shown.

## Detecting methylation in human samples

We applied our model to a biologically relevant sample with a reference methylome to confirm our observations of detectable signal differences from methylated DNA. These experiments use the emission distributions that we trained over the CpG alphabet using dataset PCR+M.SssI-timp-021216 to predict which CpGs are methylated in human samples.

We performed three sequencing runs of genomic DNA extracted from a human female lymphoblast cell line (Coriell NA12878), along with two control runs. We generated a methylation-negative control by sequencing PCR-amplified NA12878 DNA, then treated an aliquot of the PCR amplicons with M.SssI to generate a methylation-positive control. We refer to the three sequencing runs from the Coriell NA12878 as the “natural DNA” dataset, as the cell’s methylation state was preserved. When we analyze all three natural DNA runs jointly we refer to it as the “merged” data set.

We apply our trained model to calculate the log likelihood ratio between a methylated and unmethylated versions of a substring of the reference genome:

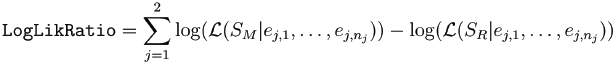

Here *S*_*R*_ is a substring of the reference genome that contains at least one CG dinucleotide (we consider nearby sites jointly as a group as each event’s current level is affected by multiple bases). *S*_*M*_ is a modified copy of *S*_*R*_ where all CG dinucleotides have been changed to MG. Presently our model cannot detect non-CG methylation, nor *k*-mers with a mix of methylated and unmethylated CGs, as our training set is limited to completely methylated or unmethylated sequence. We assume the CG sites under consideration are not hemi-methylated, e.g. both strands are methylated equally, so we sum the log-likelihood ratio from the two strands (indexed by *j*) of the 2D nanopore reads. The likelihoods are calculated by computing the probability of observing the events aligned to this portion of the reference genome, (*e*_*j,1*_,…,*e*_*j,n*_), given the hidden Markov model parameterized by *S*_*M*_ or *S*_*R*_. For complete details of this calculation see **Supplementary Note-Classifying CpG Sites**

**Figure 3** is a histogram of log likelihood ratios computed from three datasets. In this analysis we restricted the histogram to only include “singleton sites”-the CpG groups described above that only contain a single CG. The log likelihood ratio distribution for the negative control (PCR) and positive control data sets are clearly shifted to be less than zero (stronger evidence for no methylation) and greater than zero (stronger evidence for methylation), respectively. The histogram for the merged runs of natural NA12878 DNA has a single peak around zero and is wider than the distribution of the two control samples? this reflects the underlying biology as CGs for natural human DNA consist of a mixture of methylation states.

**Figure 3:**
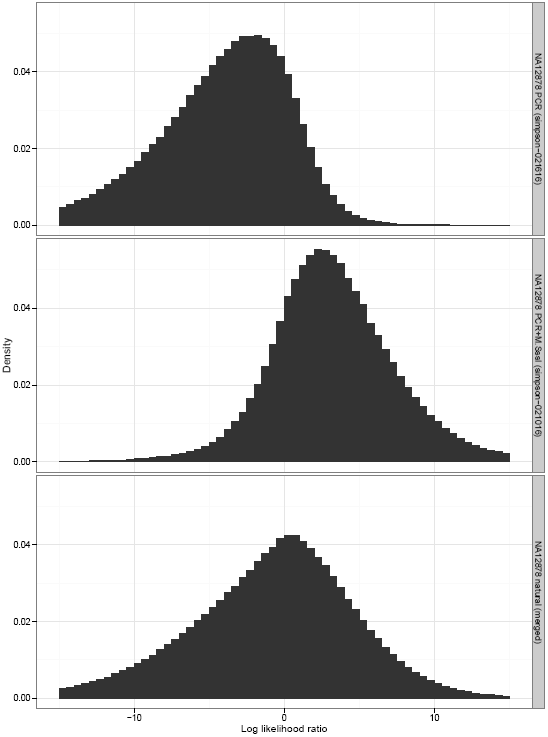
Log Likelihood Ratio comparison for human samples. A histogram of the log likelihood ratios for unmethylated NA12878 DNA (top pane), methylated NA12878 DNA (middle pane) and natural NA12878 DNA (bottom pane). In these figures we only include singleton sites.

We then assessed the accuracy of making a binary methylated/unmethylated call for singleton sites from a single 2D nanopore read. We randomly sampled 100,000 singleton sites from each of the negative and positive NA12878 control datasets then used the log likelihood ratio to make a binary classification for each site (if the log likelihood ratio was positive we classified the site as methylated). We computed accuracy by considering each site from the negative control to be unmethylated and each site from positive control to be methylated as ground truth. When considering all randomly sampled sites this classifier had 82% accuracy for singleton sites. We further characterized the accuracy of per-site methylation detection by setting a threshold on how extreme the log likelihood ratio had to be to make a call at that position (if the absolute value of the log likelihood ratio was greater than the threshold a call is made, otherwise the site is ignored). These results are shown in **Figure 4** for a threshold varying from 0 to 10. In panel A, we see that the accuracy (1-error rate) improves to over 95% as the stringency for making a call increases. However, this comes at the cost of making calls at fewer sites (panel B). When setting a threshold of 2.5 our classifier has 92% accuracy and makes calls at 62% of sites.

**Figure 4:**
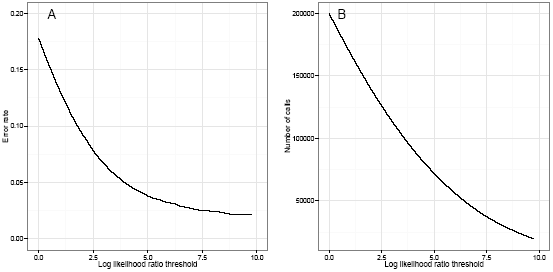
Methylation classifier error rate vs threshold. Panel A shows the error rate of the binary methylated/unmethylated classifier as a function of the log likelihood ratio threshold required to make a call. Panel B shows the number of calls made as a function of the threshold.

We also explored the biological context of our predicted methylation sites. Using a log likelihood threshold of 2.5 we made methylated/unmethylated calls for each group of sites in each sample. We calculated the percentage of called CG sites that were methylated as a function of their distance from an annotated transcription start site. We performed this analysis for the merged natural DNA dataset, the positive and negative controls and a recent bisulfite sequencing high-coverage dataset for the same sample from ENCODE (accession ENCFF279HCL). The results are shown in **Figure 5**. As expected we observe a tendency for CG sites that are near transcription start sites to be unmethylated, with percent methylation increasing as distance from the TSS increases. The pattern of methylation for the nanopore calls closely tracks the pattern for bisulfite data. This pattern is consistent across chromosomes (**Supplementary File** TSS-by-chromosome) with the exceptions of chromosome 21, which is noisy due to low coverage and few genes, and chromosome X, which has a flatter profile likely resulting from methylation arising from X-inactivation. The percentage of methylated CGs in the negative and positive controls have a low and high percent-methylated profile respectively which is independent of distance from the TSS.

**Figure 5:**
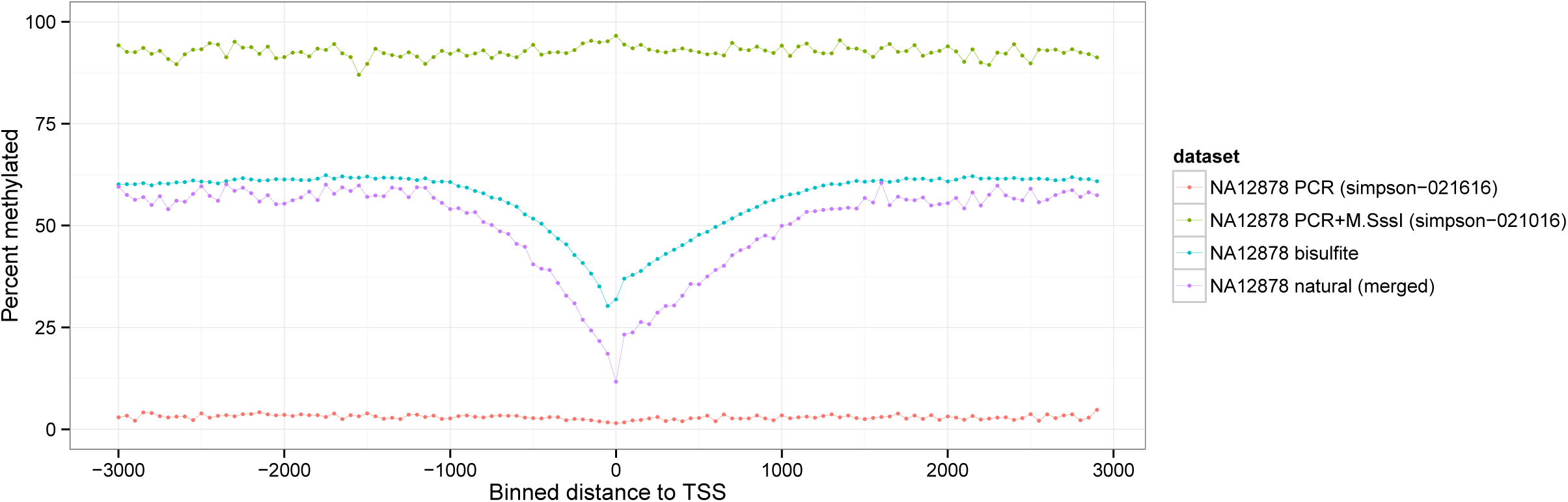
NA12878 methylation vs. distance to TSS. The percentage of CpGs predicted to be methylated as a function of their distance from an annotated transcription start site for all calls on the autosomes. The distances are binned into 50bp windows where negative distances are upstream of the TSS and positive distances are downstream, as calculated by `bedtools closest-D`.

Finally, we compared the percentage of methylation calculated by our nanopore-based method to bisulfite measurements at annotated CpG islands (CGIs). **Figure 6** show the results for the merged natural DNA data set. We find that the nanopore and bisulfite data have consistent patterns of methylation at CGIs, with overall correlation of 0.84 (Pearson’s correlation coefficient). CGIs that are in an annotated promoter show low levels of methylation, as expected. Note the quantization of the nanopore data is due to the low coverage of our data set. Repeating the calculation for our control data sets (Supplementary figures 1 and 2) breaks the correlation. In **Supplementary Figures 3**-**5** we show the three individual runs of natural NA12878 DNA to demonstrate that the detected pattern is reproducible across replicates.

**Figure 6:**
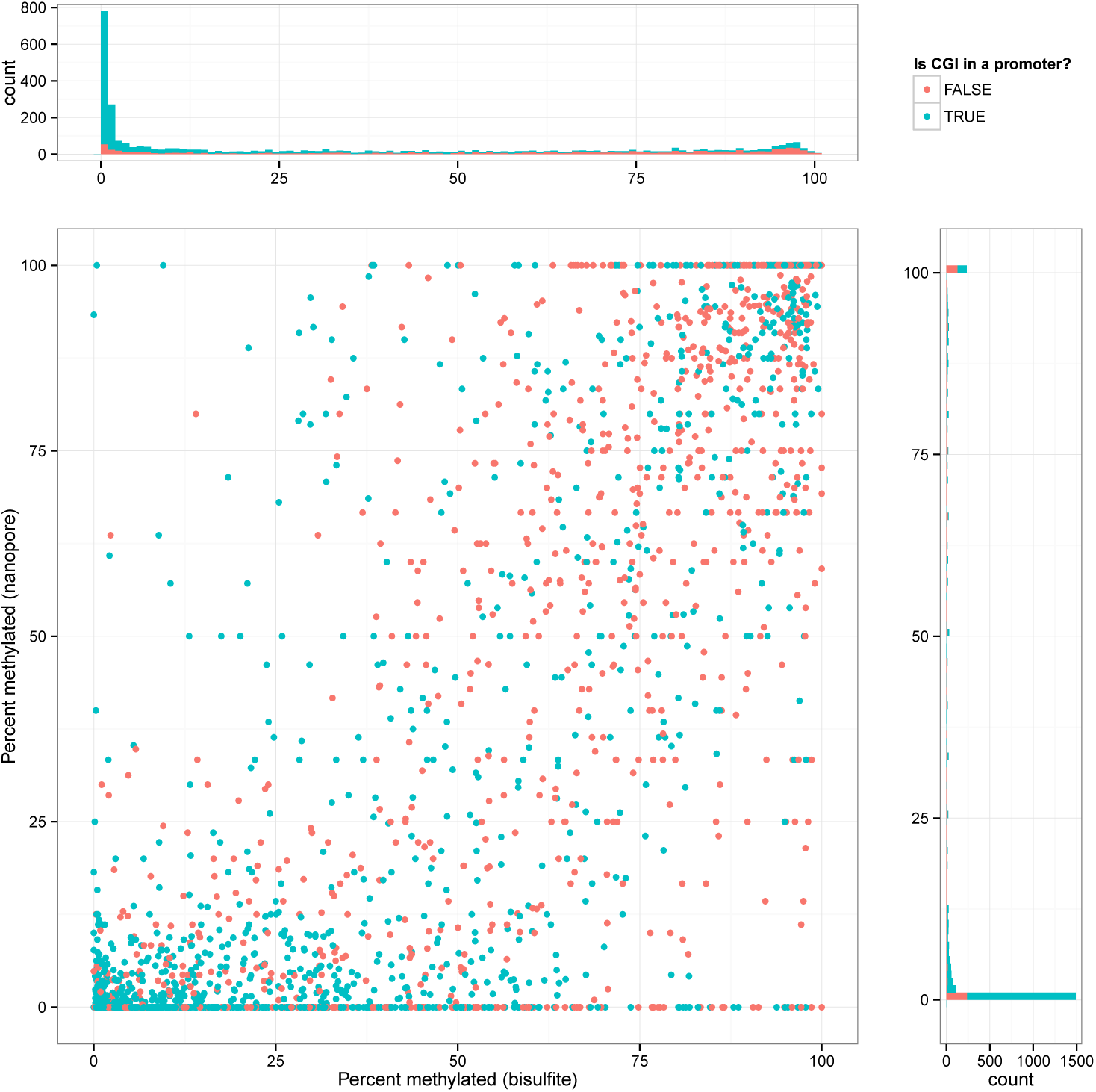
NA12878 CpG island methylation. In the main panel each point is an annotated CpG island in the human genome that was covered by both bisulfite sequencing data and nanopore reads from the merged natural NA12878 DNA data set. The x-coordinate of the point is the percentage of CpGs in the island that were predicted to be methylated from bisulfite data. The y-coordinate is the percentage of CpGs predicted to be methylated by our model using the nanopore data. The points are colored by whether the CGI is in a promoter(blue) or not(red). The histograms on the top and right of the figure are the marginal distributions of methylation percentages for the bisulfite and nanopore calls respectively.

## Discussion

In this paper we have demonstrated that 5-methylcytosine is identifiable when sequencing natural DNA using Oxford Nanopore Technologies’ MinION instrument. We calculated the accuracy of detecting isolated 5-mCs in a CpG context as 82%, increasing to 95% with more stringent calling thresholds. Our human results used low-coverage data where the interrogated sites were covered by a single nanopore read. In future work, as the throughput of the MinION and related sequencing instruments grows, we anticipate that the accuracy of our calls will improve by integrating the signals from multiple overlapping reads.

One limitation of our current model is that we used a training data set consisting of completely methylated DNA, reducing our ability to directly call heterogeneous methylation within a region. This is especially relevant in high CG density regions, so part of our next set of experiments will be the generation of more extensive training sets for all possible combinations.

There are other types of nucleotide modifications that have functional consequences in protein binding^19^ and DNA structure^20^ including 5-hydroxymethylcytosine (5-hmC), 5-formylcytosine (5-fC) and 5-carboxylcytosine (5-caC). Previous work in research pores has demonstrated discrimination of C, 5-mC, 5-hmC, 5-fC and 5-caC in a research setting? it seems likely that the general trend of detection via modulation of the nanopore current will hold true in any type of pore based sequencing^21^. We are also interested in training for adenosine variants-N6-methyladenosine has recently been shown to occur naturally in *Drosophilia*^22^ and *C. elegans* genomes^23^, even in mouse embryonic stem cells^24^. Exogenous methylation application also has applications in research and discovery of nuclear architecture or protein occupancy, e.g. damID^25^ or NOME-seq^26^. Finally, general models of other DNA damage, resulting from heavy metals, oxidation, UV damage or other alterations might be detected in natural DNA with this method - these changes are otherwise impossible to probe on a single-base level. In future work we will train models for these base modifications to generate a unified calling model that can be applied to detect interleaved patterns of these methylation marks for human DNA, allowing comprehensive epigenetic profiling from a single assay.

## Methods

We prepared and sequenced bacterial and human DNA samples under a variety of conditions using the MinION instrument. We generated data in two laboratories (Johns Hopkins University and the Ontario Institute for Cancer Research). **Supplemental Table 5** presents the metadata for each sequencing run we produced including the combination of protocols used to prepare and sequence the DNA, where the library was prepared and sequenced, how much data was generated and the ENA accession number.

### E.coli and NA12878 unmethylated, M.SssI treated and native

*Escherichia coli* K12 ER2925 unmethylated genomic DNA (Zymo Research) and genomic DNA from CEPH lymphoblastoid cell line NA12878 (Coriell Biorepository) were sheared to 8kb using Covaris G-tubes, by centrifuging 50 ng of sample in 45 ul of water at 2350x g for 1 minute, then inverting and centrifuging again with the same conditions. The sheared sample was end repaired and dA-tailed using the Ultra II end repair module (NEB), and PCR adapters (Oxford Nanopore Technologies) were ligated on using Blunt/TA Ligase Master Mix (NEB). The sample was amplified with LongAMP master mix (NEB). Half of the PCR product was set aside for sequencing (unmethylated control), and half was subjected to enzymatic methylation with M.SssI methyltransferase (CpG methylase, Zymo), which converts nearly all cytosines in a CpG context to 5-mC. The standard Zymo methylation protocol was followed, with modifications made to S-adenosylmethionine concentration (16 mM), enzyme concentration (8 units) and incubation time (18 hours). Natural *E. coli* ER2925 and NA12878 was also sheared and end repaired according to the methods described above.

### Sequencing samples on MinION

Samples were prepared for sequencing following the protocol in the genomic sequencing kit SQK-MAP006 (ONT). Approximately 1 ug of DNA prepared as described above was end repaired and dA-tailed (NEB), then subsequently cleaned with 1X Ampure XP (Beckman Coulter) and eluted in water. Adapter ligation using Blunt/TA Ligase Master Mix (NEB) affixed leader and biotinylated hairpin adapters on either end of the library molecules (SQK-MAP006 Genomic DNA Sequencing Kit, Oxford Nanopore), along with loaded motor proteins on leader adapters and tether molecules designed to bind to the synthetic membrane surface of the pore array. My-One Streptavidin C1 Dynabeads (Thermo Fisher) were used to enrich for library molecules containing the biotinylated hairpin. Libraries were eluted off streptavidin beads using elution buffer (Oxford Nanopore), added to Running Buffer and Fuel Mix, and loaded on the MinION Mk1 sequencer and run for up to 48 hrs. The PCR-treated and PCR+M.SssI treated NA12878 samples were prepared at JHU and sequenced at OICR, all other samples were prepared and sequenced within the same lab (**Supplementary Table 5**).

The *E. coli* K12 MG1655 run of PCR-amplified DNA was obtained from a public source (see accession below).

### Illumina validation

*E. coli* and reduced representation samples were assayed for CpG methylation content using bisulfite sequencing with the Illumina Miseq. 500 ng of *E. coli* DNA was sheared to 300bp using the Bioruptor Pico (Diagenode), then end-repaired and dA-tailed using NEBNext Ultra II library prep kit (NEB). Methylated adapters (NEB) were ligated on using Blunt/TA ligase Master Mix, then libraries were bisulfite converted (Zymo Methylation-Lightning). Libraries were then amplified using indexed primers following NEB recommendations, but using Kapa Hifi Uracil as the polymerase. 10 ng of reduced representation samples MDA-MB-231 and MCF10A were prepared for Illumina sequencing using Accel-NGS Methyl-Seq (Swift Biosciences). All samples were sequenced on an Illumina Miseq at JHU using v3 150 chemistry.

### Computational Methods

The details of our probabilistic model, training and classification algorithms and a description of analysis method we used to generate the results presented above are contained in the supplement. Our analysis pipeline is reproducible and documented in the GitHub repository: https://github.com/jts/methylation-analysis?. Our training and classification code is in https://github.com/jts/nanopolish?.

### Accessions

All data generated for this study was deposited in the ENA under accession PRJEB13021. In addition, we used PCR-amplified *E. coli* K12 sequencing data previously collected with accession ERR1147229.

## Author Contributions

J.T.S. and W.T designed the study. W.T., R.W. and P.C.Z. performed the experiments. J.T.S., M.D. and L.J.D. designed and implemented the methylation training and inference framework. J.T.S, W.T., R.W and P.C.Z. wrote the paper.

## Acknowledgements

The authors thank Nick Loman and Josh Quick for making the E. coli K12 dataset publicly available. We thank Andy Feinberg and Kasper Hansen for helpful discussions.

**1 Supplementary Note**

**1.1 The nanopolish hidden Markov model**

In previous work [1, 2] we developed a hidden Markov model to calculate the probability of observing a sequence of electric current *eventse*_*1*_,…,*e*_*n*_ (measured by the MinION) given a nucleotide sequence *S*. The nucleotide sequence *S* parameterizes the hidden Markov model; *S* determines the structure of the model, the transition probabilities and emission distributions used in each state.

The hidden Markov model has a block of states for every k-mer of *S* (in this work *k* = 6) with each block containing a `match` state (indicating a particular event was observed from a particular k-mer), a extra state (indicating additional observations of events from a particular k-mer, reachable from the match state) and a `skip` state (indicating no event observations for a k-mer). In addition we have a `softclip` state that allows events at the start and end of the event sequence to go unaligned, to account for uncertainty in the end points of the event sequence.

Throughout this text we will refer to the likelihood of a sequence *S*, which is defined to be the probability of observing the sequence of events given the hidden Markov model parameterized by *S*:

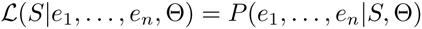

Here Θ is the complete set of model parameters; the transition distributions are as previously described [1]. In the next section we describe the emission distributions.

**1.1.1 Emission distributions**

We follow the Oxford Nanopore Technologies basecaller (Metrichor) by modeling the probability of observing an event ei given that the true sequence in the pore is k-mer *k* using a Gaussian distribution:

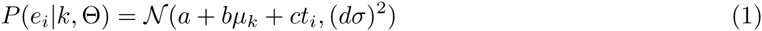

where:

- *e*_*i*_: the measured current level for event *i*
- *t*_*i*_: the time at which event *i* is observed
- *k*: the *k*-mer
- *µ*_*k*_: the mean current level for k-mer *k*
- σ_*k*_: the standard deviation of the current level for k-mer *k*
- *a*: the Metrichor `shift` parameter
- *b*: the Metrichor `scale` parameter
- *c*: the Metrichor `drift` parameter
- *d*: the Metrichor `var` parameter

The Metrichor parameters *a, b, c, d* are used to capture read-specific deviations from the base model 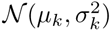. These values are provided within the HDF5-formatted FAST5 files. In this work we use these values without modification.

Equation 1 is the emission distribution for the `match` states of the HMM where *k* is the k-mer corresponding to the state. The emission distribution for the `extra` states has a similar form but we scale the standard deviation by an additional factor *v* to account for more noise in the subsequent observations from a k-mer (refer to [1] for more details). The `skip` state does not emit observations. The `softclip` state has a log-scaled emission probability of-3.0 independent of the event observation.

When base-calling a read, Metrichor selects a set of emission distributions from one of three models, depending on the inferred physical properties of the sequenced DNA. These models are called “template”, “complement.pop1” and “complement.pop2”. The template model is always used for the first sequencing strand; if a hairpin is detected, one of the two complement models is selected for the second strand. Within our hidden Markov model we always use the same strand model selected by Metrichor. When training new emission distributions later we will train the three strand models separately.

It is our goal to calculate whether a particular reference base is methylated. To do this we need to expand upon the provided ONT models to include *k*-mers that contain methylated bases.

**1.1.2 Expanding the emission alphabet**

Previously our hidden Markov model used the four base nucleotide alphabet Σ_nuc_ = {`A, C, G, T`}. The ONT-provided strand models therefore consisted of 4^6^ = 4096 different Gaussian distributions.

To handle methylation we expand the alphabet to include a new symbol M representing 5-methylcytosine (Σ _CpG_ = {`A, C, G, T, M`}). This increases the number of emission distributions to 5^6^ = 15625, but as we are only interested in measuring 5mC in a CpG context in this work, many of these *k*-mers are invalid methylation sites. For example `AMTAGA` is an invalid k-mer in this model as the methylated cytosine is not followed by a G but `GTAMGA` and `ACGATM` are valid *k*-mers. In the main text we refer to this as the CpG alphabet.

Later we will refer to the unmethylated version of a k-mer. By this we mean changing all of the M symbols to C. For example AACGAA is the unmethylated version of `AAMGAA`.

**1.2 Training Emission Distributions**

In this section we describe how we use MinION reads aligned to a known reference genome to learn new parameters for the emission distributions for each k-mer for each of the three ONT strand models.

First we extract the “two-direction” (2D) reads from a MinION run using `poretools` [3]. **In** this study we use both “pass” and “fail” reads as reads containing methylated CpGs may have lower quality scores. We then use `bwa mem-x ont2d` [4] to align these reads to the reference genome. Reads that were ambiguously mapped (mapping quality 0) were discarded. We iterate over the 2D reads aligned in base-space and realign them in event-space using the method described below in section 1.2.1. During this procedure we collect a list of events aligned to each k-mer for each ONT strand model. An event is used if it is not in the first or last 5 events of the alignment, if it was aligned to a match state of the HMM and if its duration is at least 5 milliseconds. After realigning all reads we use the procedure described in section 1.2.2 to learn new parameters 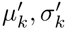 for all *k*-mers with at least 100 aligned events. After new parameters are learned we start again from the realignment step for a total of five iterations.

We use the region from 50,000 bp to 3, 250,000 bp of the *E. coli* K12 MG1655 reference genome for training over the nucleotide alphabet. We initialize the Gaussian parameters to the values in the ONT models.

Prior to training *k*-mers over the CpG alphabet we change all `CG` dinucleotides in the reference to `MG`. We use the same training region as above. The initial Gaussian parameters for unmethylated *k*-mers are taken from the ONT models. For methylated *k*-mers the parameters are initialized to the parameters for the unmethylated version of the k-mer.

**1.2.1 Aligning Events to *k*-mers**

We use the Viterbi algorithm [5] on our HMM to obtain an alignment between events and *k*-mers of *S*. When *S* is a known substring of a reference genome, we therefore obtain an alignment between events measured by the MinION and reference *k*-mers.

To reduce compute time we align events to the reference genome in segments of 50 events using the base-space alignments as a guide. We progressively align each segment using the aligned coordinates of the previous segment to calculate the start point. This progressive event alignment algorithm is implemented in the function `align_read_to_ref` in `nanopolish_eventalign.cpp`.

**1.2.2 Fitting a Gaussian Mixture Model**

Let 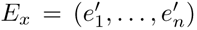 be the events aligned to k-mer *x* where 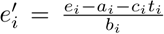 is a transformation of ei to account for the Metrichor shift, scale and drift parameters. To account for incomplete enzymatic methylation in the training data set, we fit a two-component Gaussian mixture model to *E*_*x*_ when *x* contains a methylated base (eg *x* has an `M` symbol):

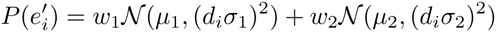

Here *w*_1_, *w*_2_ are mixture weights representing the proportion of events that originate from methylated and unmethylated sequence, respectively (*w*_1_+*w*_2_ = 1). Similarly, *µ*_1_, *σ*_1_, *µ*_2_, *σ*_2_ are Gaussian parameters for methylated and unmethylated versions of *x*.

We fit *w*_*j*_, *µ*_j_ *σ*_j_ using ten iterations of the expectation-maximization algorithm. Let *Zi* be a random variable indicating the component observation *i* comes from and *x*_*j*_ be the k-mer for the *j*-th component of the mixture. First we calculate the *responsibility* of component *j* for observation *i*:

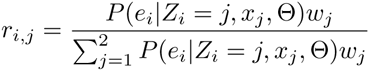

We update the weights and Gaussian parameters as follows:

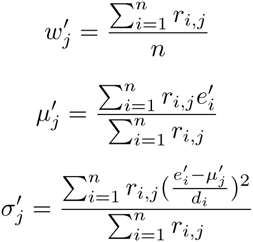

When training *k*-mers with a methylation site we initialize *w*_1_ = 0.95 and set the parameters for the k-mer to be 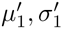, from the final iteration. When training *k*-mers that do not contain methylation sites we simply fit a Gaussian distribution using equations similar to the 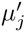 and 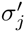 updates above.

**1.2.3 Training Data Sets**

In the main text we describe how we generated two types of training data - one type is PCR amplified DNA, to remove any methylation and DNA damage, and the other type is PCR amplified DNA treated with M.SssI, which will methylate nearly all cytosines in a CpG context. We train using both types of data over the both the nucleotide and CpG alphabet, generating four combinations of data type/alphabet (as shown in **Supplemental Tables 1**-**4** and discussed in the results of the main text). We note however that not all *k*-mers over the CpG alphabet can be trained. As we change all CG dinucleotides in the reference genome to MG, there will be no events aligned to *k*-mers containing a CG and these will go untrained, defaulting to the ONT-provided values. Importantly *k*-mers with a mixture of methylation sites (like `ACGMGT`) cannot be trained. This limitation has an important effect on calling methylation, which we discuss in 1.3.

**1.2.4 Assessing Model Training**

The output of our training method is a new set of parameters 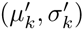 for every trained k-mer in the alphabet, for each of the three strand models. To assess how well our training step performs we compared the trained parameters to the ONT-provided parameters (*µ*_*k*_,*σ*_*k*_). We counted the number of *k*-mers where the difference in means was greater than a threshold, 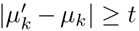 for *t* ∈ (0.1, 0.5,1.0, 2.0). This analysis code is in `generate_training_table.py`.

**1.3 Classifying CpG sites**

**In** this section we describe how we call CpG sites for sequenced human genomes using our trained model. First we describe the terminology we will use and give a high-level overview of the algorithm. In the subsequent sections we describe the individual steps in more detail.

We will refer to a substring of the reference genome that we are testing for methylation as *S*_*R*_. Below, we will construct *S_R_* such that it contains at least one, and possibly multiple, `CG` nucleotides plus flanking sequence that does not contain a `CG`. We will refer to the `CG`s within *S_R_* as a *group* of sites. We group sites together as the measured event current levels do not correspond to a single base but rather a k-mer. For this reason we must jointly test closely-separated `CG`s for methylation in the same manner that we tested haplotypes containing combinations of SNPs in previous work [2]. However, a limitation of our training data is that it consists of completely methylated genomic DNA and we are unable to train *k*-mers with a mixture of methylation patterns (for example we cannot train the k-mer `ACGTMG`). When testing for methylation we assume all sites in the group have the same methylation status. We refer to the methylated version of *S_R_* as *SM*. For example if *S_R_* = AAAACGCGAAAAA then *SM* = AAAAAMGMGAAAAA. This assumption is reasonable as it is expected that the methylation status of nearby sites is correlated [6, 7]. In future work we aim to remove this assumption. Often *S_R_* will contain only a single `CG` dinucleotide. We call these cases “singleton sites” and for some analyses later we will only consider such sites.

Our procedure for classifying CpG sites as methylated or not has two main steps. First we align basecalled nanopore reads to a reference genome and use the alignments to find and group the `CG` sites that are covered by a read. We then call each group as methylated or unmethylated using log likelihood ratios. We describe the two main components of the algorithm separately in the following sections.

**1.3.1 Alignment and CpG site discovery**

We begin by extracting 2D nanopore reads (pass and fail) for the run using `poretools`. We align the extracted reads to the human reference genome (build GRCh38.p5 downloaded from Gencode) using `bwa mem-x ont2d`. Reads that are ambiguously mapped (mapping quality 0) are discarded. For each remaining aligned read we iterate over the aligned segment of the reference genome and extract the locations of the `CG` dinucleotides. Let *c*_1_,…, *c*_*n*_ be the reference coordinates of the extracted `CG` sites for the current read. We partition the list of covered CG sites into *groups* that are separated by at least 10bp. Each group is tested for methylation independently. Let *c*_*s*_ and *c*_*e*_ be the position of the first and last `CG` in the current group. We extract the reference substring, *S*_*R*_, along with 10bp of flanking sequence: *S*_*R*_ = `reference`[*c*_*s*_ — 10: *c*_*e*_ + 10]. We also extract from the aligned nanopore reads the template and complement events aligned within the reference range [*c*_*s*_ — 10,…, *c*_*e*_ + 10]. We calculate a log likelihood ratio between *S*_*M*_ and *S*_*R*_ using the method described 1.3.2 for the template and complement strands independently. We assume the sequenced DNA is not hemimethylated so the log likelihood ratio for the group is the sum of the log likelihood ratio for the two independent strands. This ratio, along with the reference coordinates of the group, the number of CpG sites in the group, and *S_R_* is output in a BED-formatted file for downstream analysis.

**1.3.2 Calculating likelihood ratios**

**In** the previous section we described how we extract *S*_*R*_, a substring of a reference genome that contains one or more `CG` sites and at least 10bp for flanking sequence that does not contain a `CG`. *S_R_*, along with a sequence of events *e*_*j,1*_,…, *e*_*j,n*_j__ from sequencing strand *j* (either template or complement) of a nanopore read is the input into the likelihood ratio calculation. Recall that *S*_*M*_ is defined to be a modified version of *S_R_* where all `CG` dinucleotides are converted to `MG`.

In section 1.1 we defined the likelihood of an arbitrary sequence S to be the probability of observing the sequence of nanopore events given a hidden Markov model parameterized by S. To calculate the likelihoods used in this section we use the same method however we now use the trained emission distributions over the Σ_CpG_ alphabet for both *S*_*M*_ and *S*_*R*_. Thus, the likelihood ratio between the completely-methylated and unmethylated reference sequence is:

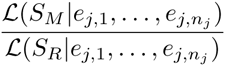

The log likelihood ratio used in section 1.3.1 is simply the log of this value, summed over the two strands:

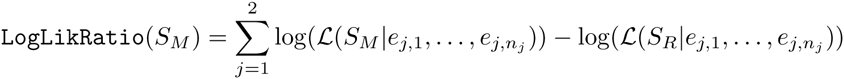

**1.3.3 Classification**

In some analyses we will make a binary methylated or unmethylated classification for groups of `CG` sites rather than working directly with the log likelihood ratio. To improve the accuracy of this binary classification we set a threshold t on how extreme the log likelihood ratio must be to make a call. For a group consisting of n `CG` sites, we only make a call if |`LogLikRatio`(S_M_)| > nt otherwise the group is ignored. When calling a group, we say that n sites are called and of the called sites n are methylated if `LogLikRatio`(*S*_*M*_) > 0.

**1.3.4 Source code**

The complete source code for the methods described in this section can be found in function `calculate methylation for read` in `nanopolish methyltest.cpp`.

**1.4 Evaluating Accuracy**

To evaluate the accuracy of our methylation predictions we randomly sampled 100, 000 singleton sites (groups containing only one `CG`) from each of the unmethylated and methylated NA12878 control samples. We then calculated the number of these sampled sites that were correctly identified as methylated or unmethyated, where the truth for each site was defined to be the data set that site originated from (all sites sampled from the methylated data set were considered to be truly methylated and vice-verse). We performed this analysis for call thresholds t ranging from 0 to 10. In Figure 4 of the main text we plot the error rate of this classifier and the total number of calls as a function of t. This analysis code is in `calculate call accuracy.py`.

**1.5 Transcription Start Site Analysis**

To calculate the percentage of CpGs that were methylated as a function of the distance to a transcription start site (TSS) we generated a new database of TSSs using the script https://github.com/sdjebali/MakeGencodeTSS on the Gencode v24 release (http://www.gencodegenes.org/releases/24.html). This procedure is documented here:

https://github.com/jts/methylation-analysis/blob/master/annotations/README

We then used `bedtools closest-D` [8] to calculate the distance between each call and a TSS for both the bisulfite data and the three nanopore data sets (NA12878 natural DNA and the two controls). We parsed the resulting annotated BED files and counted the total number of CpG sites covered by a read and the number of those sites called as methylated in 50bp bins in the range-3000bp to 3000bp. Here negative distances refer to sites that are upstream of the TSS and positive distances are downstream of the TSS. For the nanopore data we set the call threshold at 2.5. When a called group in the nanopore data set had multiple CpG sites, they were counted together. For example if a group had 3 CpG sites and the group was called as methylated, we counted 3 methylated calls in the corresponding bin.

In **Figure 5** in the main text we plot the aggregated results for the autosomes. In Supplementary File TSS-by-chromosome we plot the results chromosome-by-chromosome.

The analysis code is in `calculate methylation by distance.py`

**1.6 CpG Island Analysis**

We downloaded a database of CpG islands (CGI) for the GRCh38 reference genome from the UCSC genome browser. We generated a BED file of promoters regions defined as 2000bp upstream and 200bp downstream of the TSSs described above. We used `bedtools map` to annotate whether each CGI was within a promoter. We then used `bedtools intersect` to find all methylation calls that overlap an annotated CGI for the bisulfite data and all nanopore data sets. For each CGI we calculate the percentage of methylated CpGs by counting the total number of CpGs in the CGI that were covered by a read and counting the number of those CpGs that were classified as methylated. This is implemented in `calculate_methylation_at_cpg islands.py`. In the output figures we plot the bisulfite percent methylated against one of the nanopore datasets for CGIs that were covered by reads in both datasets.

**1.7 Reproducibility**

The analysis pipeline is implemented as a Makefile that begins from the raw data downloaded from the ENA, downloads dependencies and versioned analysis code, runs our training method, calls methylation for the human nanopore runs and generates the tables and figures in this paper. This Makefile, which is annotated with a description of each step, is located on github:

https://github.com/jts/methylation-analysis/blob/master/pipeline.make

---

**Table 1:** Model training results for PCR-treated DNA over the nucleotide alphabet. The final four fields are the number of *k*-mers where the mean of the trained Gaussian differs from the ONT-trained mean by more than x pA.

| model | sample | run | training events | trained kmers | 0.1 | 0.5 | 1.0 | 2.0 |
| --- | --- | --- | --- | --- | --- | --- | --- | --- |
| template | *E. coli* ER2925 (PCR) | timp 113015 | 70M | 4096 | 3257 | 589 | 16 | 0 |
|  | *E. coli* ER2925 (PCR) | timp 021216 | 123M | 4096 | 2261 | 72 | 3 | 0 |
|  | *E. coli* K12 (PCR) | loman 250915 | 76M | 4096 | 2149 | 50 | 2 | 0 |
| comp.pop1 | *E. coli* ER2925 (PCR) | timp 113015 | 23M | 4091 | 3041 | 417 | 21 | 0 |
|  | *E. coli* ER2925 (PCR) | timp 021216 | 25M | 4091 | 2613 | 219 | 19 | 0 |
|  | *E. coli* K12 (PCR) | loman 250915 | 14M | 4086 | 2551 | 221 | 23 | 0 |
| comp.pop2 | *E. coli* ER2925 (PCR) | timp 113015 | 44M | 4095 | 3392 | 1305 | 283 | 4 |
|  | *E. coli* ER2925 (PCR) | timp 021216 | 89M | 4096 | 2497 | 224 | 21 | 1 |
|  | *E. coli* K12 (PCR) | loman 250915 | 56M | 4096 | 2095 | 77 | 9 | 0 |

**Table 2:**
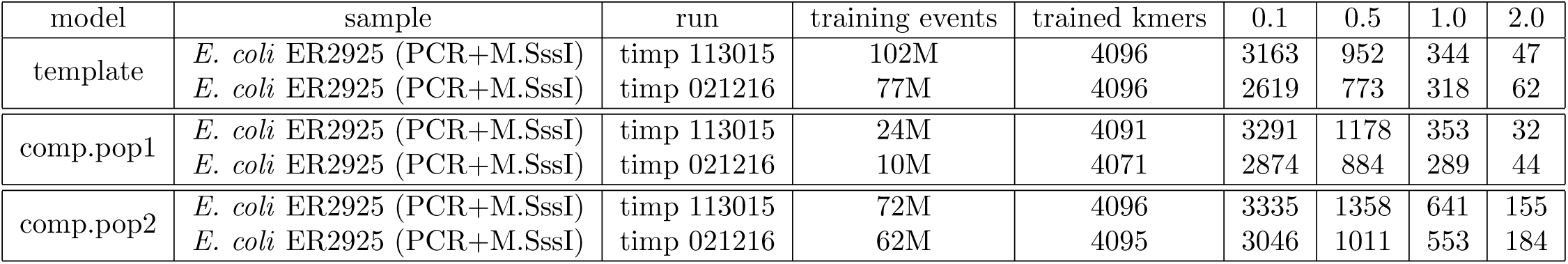
Model training results for PCR+M.SssI-treated DNA over the nucleotide alphabet. The final four fields are the number of *k*-mers where the mean of the trained Gaussian differs from the ONT-trained mean by more than x pA.

**Table 3:** Model training results for PCR-treated DNA over the cpg alphabet. The final four fields are the number of *k*-mers where the mean of the trained Gaussian differs from the ONT-trained mean by more than x pA.

| model | sample | run | training events | trained kmers | 0.1 | 0.5 | 1.0 | 2.0 |
| --- | --- | --- | --- | --- | --- | --- | --- | --- |
| template | *E. coli* ER2925 (PCR) | timp 113015 | 70M | 5120 | 4077 | 867 | 76 | 38 |
|  | *E. coli* ER2925 (PCR) | timp 021216 | 123M | 5120 | 2972 | 158 | 18 | 6 |
|  | *E. coli* K12 (PCR) | loman 250915 | 76M | 5120 | 2781 | 110 | 16 | 9 |
| comp.pop1 | *E. coli* ER2925 (PCR) | timp 113015 | 23M | 5107 | 3873 | 632 | 80 | 11 |
|  | *E. coli* ER2925 (PCR) | timp 021216 | 25M | 5107 | 3390 | 468 | 76 | 1 |
|  | *E. coli* K12 (PCR) | loman 250915 | 14M | 5097 | 3333 | 351 | 48 | 5 |
| comp.pop2 | *E. coli* ER2925 (PCR) | timp 113015 | 44M | 5117 | 4240 | 1650 | 458 | 33 |
|  | *E. coli* ER2925 (PCR) | timp 021216 | 89M | 5119 | 3251 | 417 | 71 | 5 |
|  | *E. coli* K12 (PCR) | loman 250915 | 56M | 5119 | 2872 | 180 | 33 | 2 |

**Table 4:** Model training results for PCR+M.SssI-treated DNA over the cpg alphabet. The final four fields are the number of *k*-mers where the mean of the trained Gaussian differs from the ONT-trained mean by more than x pA.

| model | sample | run | training events | trained kmers | 0.1 | 0.5 | 1.0 | 2.0 |
| --- | --- | --- | --- | --- | --- | --- | --- | --- |
| template | *E. coli* ER2925 (PCR+M.SssI) | timp 113015 | 101M | 5120 | 4006 | 1683 | 997 | 331 |
|  | *E. coli* ER2925 (PCR+M.SssI) | timp 021216 | 77M | 5120 | 3339 | 1554 | 1038 | 303 |
| comp.pop1 | *E. coli* ER2925 (PCR+M.SssI) | timp 113015 | 24M | 5108 | 4248 | 1900 | 971 | 307 |
|  | *E. coli* ER2925 (PCR+M.SssI) | timp 021216 | 10M | 5068 | 3743 | 1631 | 960 | 320 |
| comp.pop2 | *E. coli* ER2925 (PCR+M.SssI) | timp 113015 | 72M | 5120 | 4255 | 2028 | 935 | 302 |
|  | *E. coli* ER2925 (PCR+M.SssI) | timp 021216 | 62M | 5120 | 3874 | 1665 | 927 | 300 |

**Table 5:** Metadata for sequencing experiments generated for this study

| structured name | library/sequencing lab | library protocols | 2D yield | accession |
| --- | --- | --- | --- | --- |
| ecoli er2925.MSssI.timp.100215 | Timp/Timp | MSssI + ONT gDNA sequencing kit | 54.8 Mbp | ERR1309542 |
| ecoli er2925.MSssI.timp.100615 | Timp/Timp | MSssI + ONT gDNA sequencing kit | 89.5 Mbp | ERR1309543 |
| ecoli er2925.native.timp.102615 | Timp/Timp | ONT gDNA sequencing kit | 94.6 Mbp | ERR1309544 |
| ecoli er2925.native.timp.110915 | Timp/Timp | ONT gDNA sequencing kit | 107.1 Mbp | ERR1309545 |
| ecoli er2925.pcr MSssI.timp.113015 | Timp/Timp | ONT gDNA low input (SssI modification) | 243.8 Mbp | ERR1309549 |
| ecoli er2925.pcr.timp.113015 | Timp/Timp | ONT gDNA low input | 165.4 Mbp | ERR1309547 |
| NA12878.native.simpson.101515 | Simpson/Simpson | ONT gDNA sequencing kit | 121.6 Mbp | ERR1309550 |
| NA12878.native.simpson.103015 | Simpson/Simpson | ONT gDNA sequencing kit | 88.9 Mbp | ERR1309551 |
| NA12878.native.timp.093015 | Timp/Timp | ONT gDNA sequencing kit | 92.2 Mbp | ERR1309552 |
| NA12878.pcr MSssI.simpson.021016 | Timp/Simpson | ONT gDNA low input (SssI modification) | 80.2 Mbp | ERR1309554 |
| NA12878.pcr.simpson.021616 | Timp/Simpson | ONT gDNA low input (Timp) | 177.8 Mbp | ERR1309553 |
| ecoli er2925.pcr.timp.021216 | Timp/Timp | ONT gDNA low input | 309.5 Mbp | ERR1309546 |
| ecoli er2925.pcr MSssI.timp.021216 | Timp/Timp | ONT gDNA low input (SssI modification) | 211.3 Mbp | ERR1309548 |

**Figure 1:**
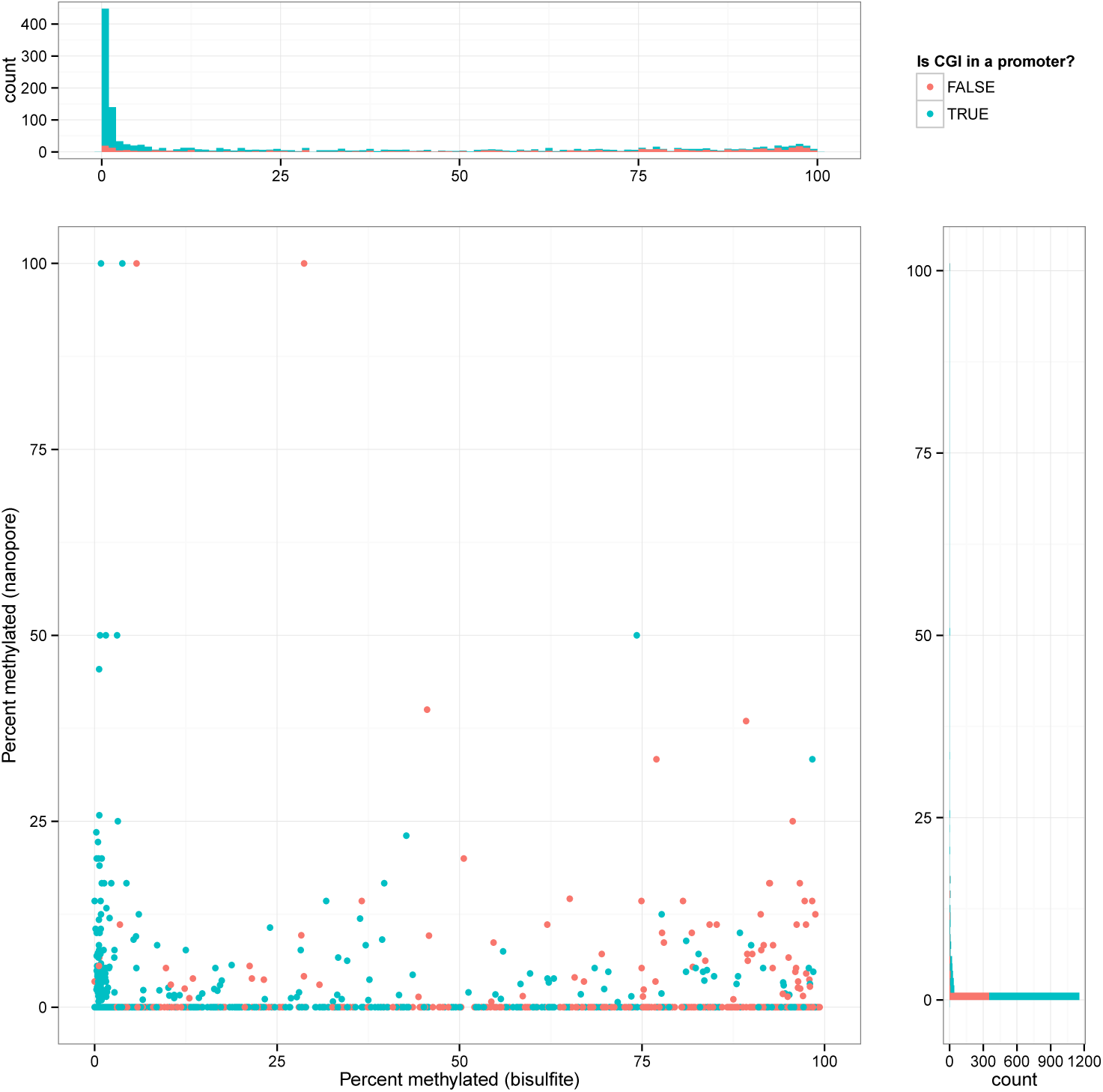
Bisulfite vs nanopore CGI methylation for the dataset PCR-simpson-021616

**Figure 2:**
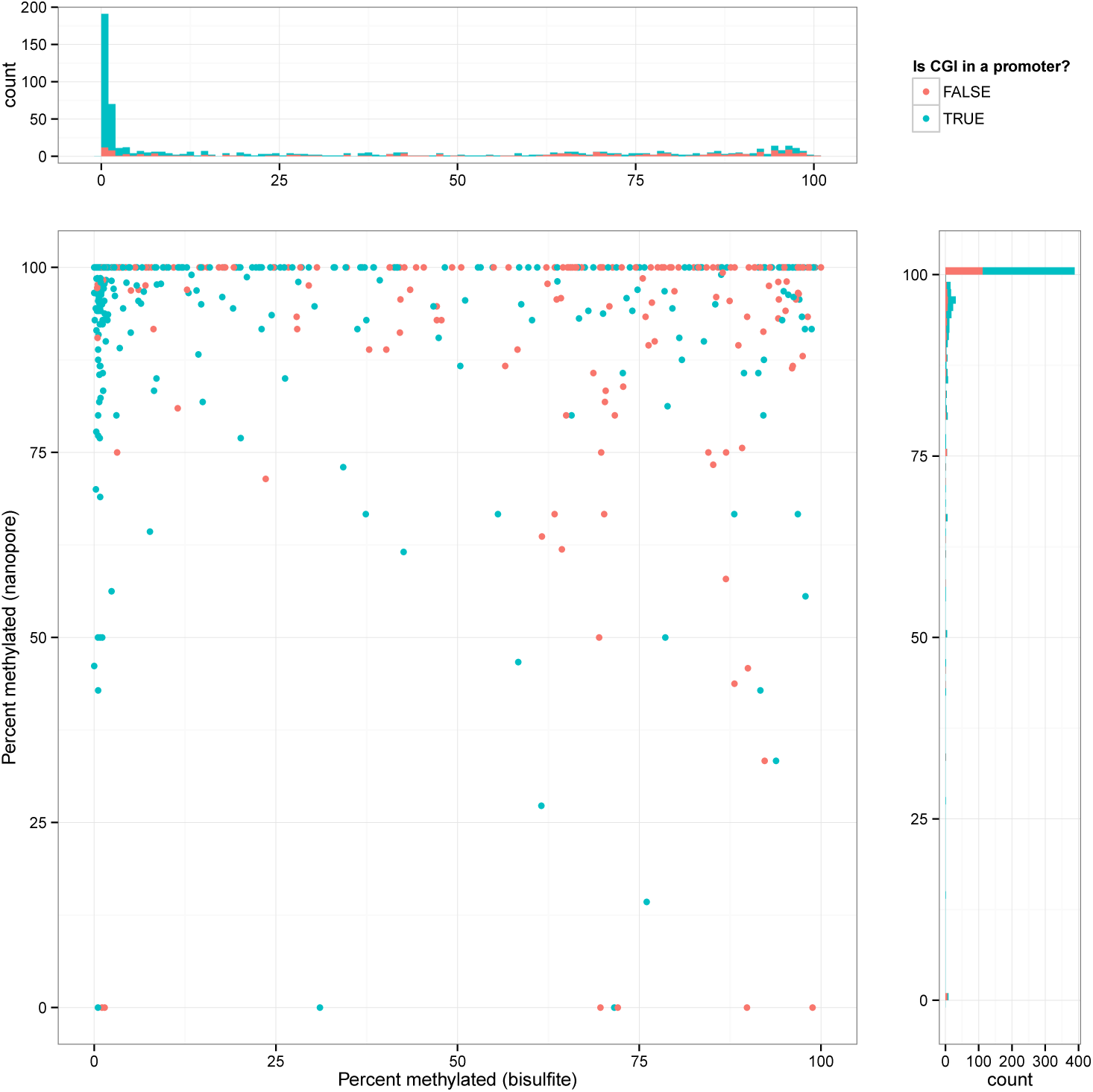
Bisulfite vs nanopore CGI methylation for the dataset PCR+M.SssI-simpson-021016

**Figure 3:**
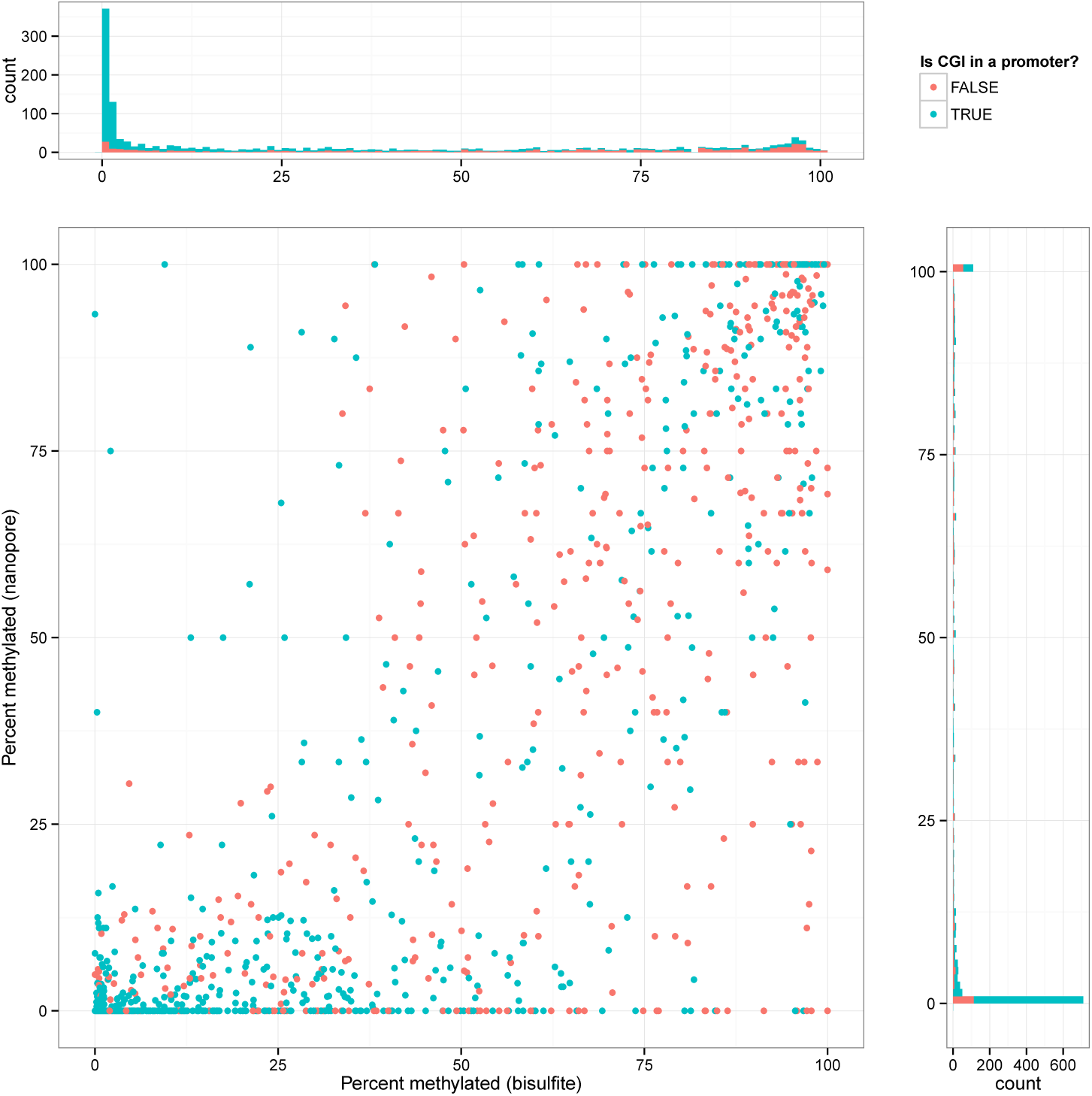
Bisulfite vs nanopore CGI methylation for the dataset natural-timp-093015

**Figure 4:**
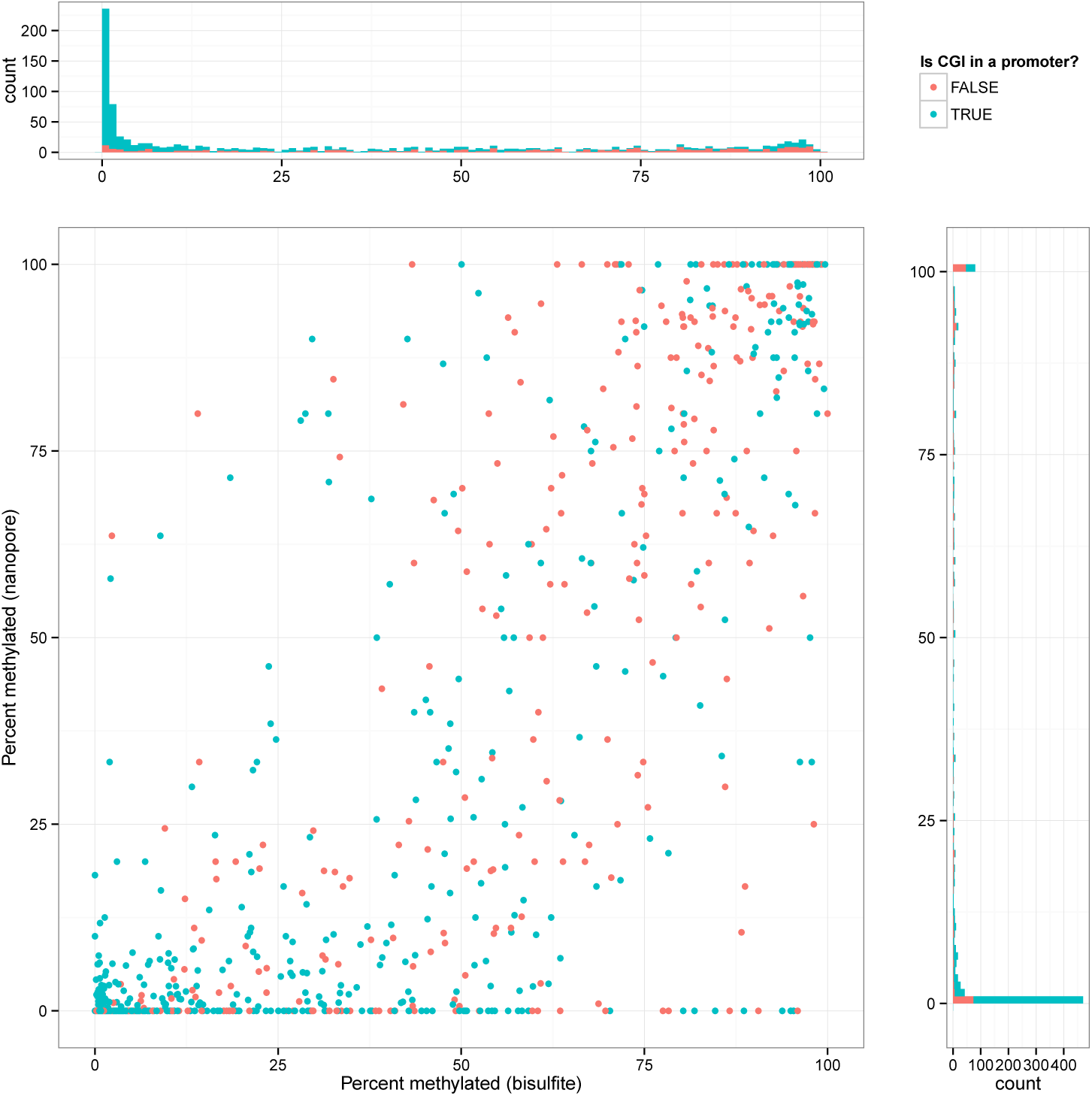
Bisulfite vs nanopore CGI methylation for the dataset natural-simpson-101515

**Figure 5:**
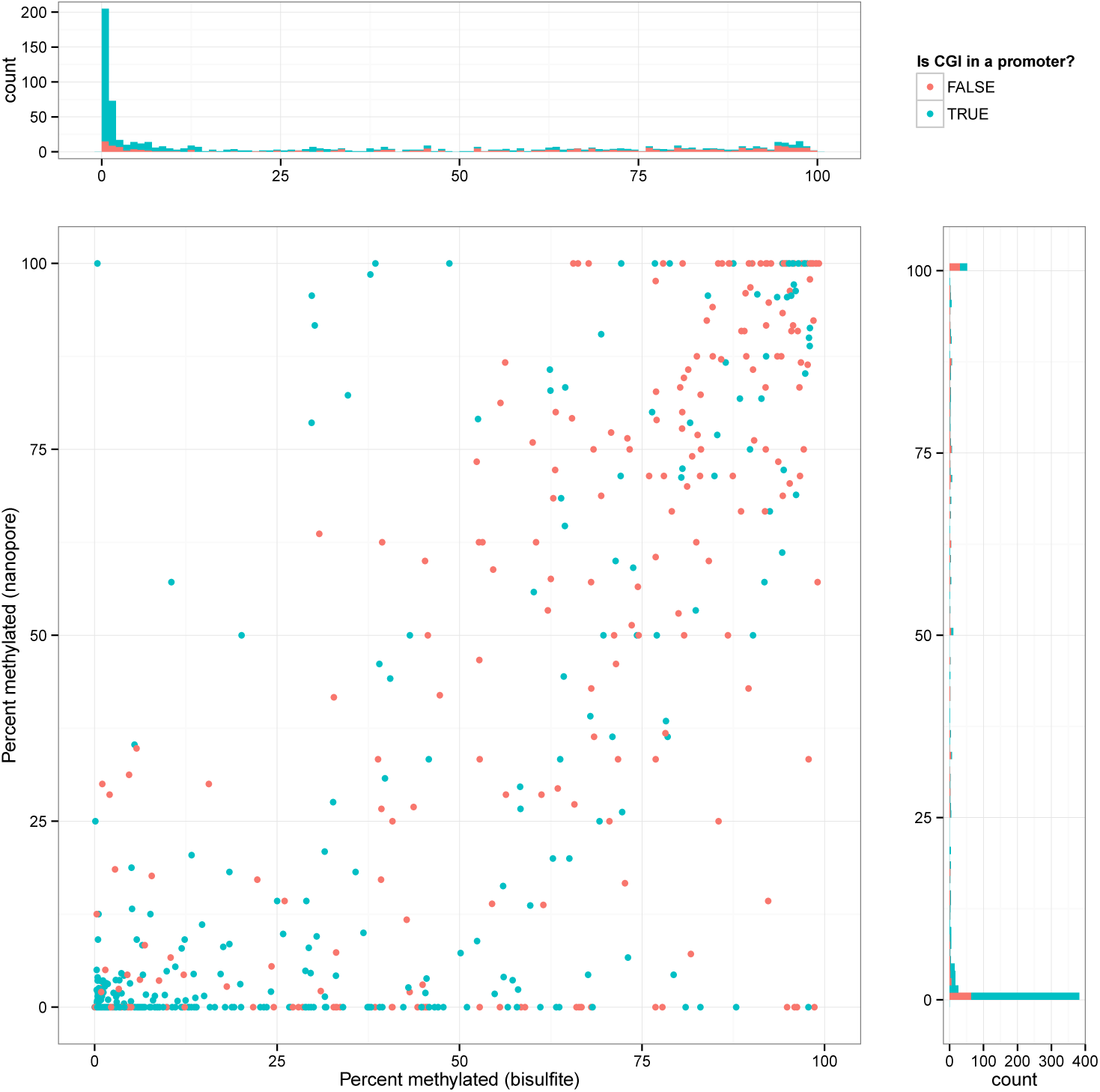
Bisulfite vs nanopore CGI methylation for the dataset natural-simpson-103015

